# Analysis of live cell data with G-DNABERT supports a role for G-quadruplexes in chromatin looping

**DOI:** 10.1101/2024.06.21.599985

**Authors:** Dmitry Konovalov, Dmitry Umerenkov, Alan Herbert, Maria Poptsova

## Abstract

Alternative DNA conformation formed by sequences called flipons potentially alter the readout of genetic information by directing the shape-specific assembly of complexes on DNA The biological roles of G-quadruplexes formed by motifs rich in guanosine repeats have been investigated experimentally using many different methodologies including G4-seq, G4 ChIP-seq, permanganate nuclease footprinting (KEx), KAS-seq, CUT&Tag with varying degrees of overlap between the results. Here we trained large language model DNABERT on existing data generated by KEx, a rapid chemical footprinting technique performed on live, intact cells using potassium permanganate. The snapshot of flipon state when combined with results from other in vitro methods that are performed on permeabilized cells, allows a high confidence mapping of G-flipons to proximal enhancer and promoter sequences. Using G4-DNABERT predictions,with data from ENdb, Zoonomia cCREs and single cell G4 CUT&Tag experiments, we found support for a model where G4-quadruplexes regulate gene expression through chromatin loop formation.

## Introduction

### G4 flipons

Sequence searches based on the (G_3-5_N_1-7_)_4_ motif revealed that the G4-quadruplexes are enriched in gene promoters. Experimental approaches based on genome wide ENCODE ChIP-seq experiments showed that these G4 motifs overlap binding sites for hundreds of transcription factors (TFs), although the resolution of such studies is limited to 150-200 bases ^1^ and some reagents like the G4-specific BG4 antibody construct detect only parallel stranded G4 structures instead of the many other G4 conformations possible ^2^. Analysis of almost 400 PDB structures revealed that there are over 48 unique quadruplex motifs, each with different configurations of the propeller, bulge, diagonal, and lateral loops that connect the quartets ^3^. One extensively investigated G4-flipon is present in the P1 promoter of the MYC gene, whose protein product plays important roles in promoting various cancers. Indeed, G4 formation enhances the engagement of various TFs that increase the transcription rate from the MYC P1 promoter, regardless of whether of which strand forms G4. Further, transcription increases in parallel with increasing G4 stability ^4^.

Overall, G4s in cells are formed transiently, with helicases able to dismantle them while other factors stabilize their formation. Experimental proofs of the reversibility of G4 formation include the use of fluorescently labeled reagents that have high specificity for quadruplexes ^5^ that reveal the resolution of quadruplexes formed during cellular stress once cells are returned to normal conditions ^6^. The G4 flip back to B-DNA can occur in minutes and may explain why chemical mapping methods based on exposure to dimethyl sulfate for many minutes previously failed to detect their presence in cells as such reagents prevent G4 reformation by modifying unfolded ssDNA ^7^. In contrast, prolonged exposure to reagents that stabilize G4s can induce formation of G4s not usually found in cells and increase rates of replication- and transcription-dependent DNA damage ^8^.

### Genome-wide experimental detection methods

There have been many techniques developed for the genome-wide detection of G4 structures. One is G4-seq, which detects DNA polymerase stalling at quadruplexes formed during sequencing ^9^. G4-seq detects more than 700 000 G4s. The set contains many noncanonical G4s with longer loops, and those with two tetrads and with bulges. ChIP-seq methods can be divided into approached using antibodies to proteins that show high affinity for G4s and antibodies that bind directly to quadruplexes. G4 ChiP-seq (antibodies to G4) identified around 10,000 G4s in nucleosome depleted regions, but are limited by the BG4 antibody used that has a preference for parallel stranded quadruplexes ^2, 10^. G4s from this set are enriched in the promoters and 5′ UTRs of highly transcribed genes, particularly in genes related to cancer and in somatic copy number amplifications. Permanganate/S1 nuclease footprinting ^11^, performed in intact cells with a 70 second exposure to the chemical, identified single-stranded regions harboring different non-B DNA structures, including G4-quadruplexes formed in around 53,000 regions.

Similar to Permanganate/S1 nuclease footprinting, N3-kethoxal–assisted labeling, called KAS-seq, is based on single-stranded DNA (ssDNA) profiles ^12^. This method exposes intact cells to chemicals for less than 10 minutes and identified around 36,000 G4s. In contrast, the CUT&Tag (cleavage under targets and tagmentation) method is based on permeabilized nuclei and long incubation periods with the BG4 antibody-tethered Tn5 tagmentation reagent. G4 CUT&Tag method detects around 18 000 G4s ^13^.

### Computational methods

Regular classic pattern detects over 370,000 G4 sequence motifs in the human genome depending on the motif searched for ^14^. G4s detected by a regular pattern are enriched in telomeres, promoters and 5’UTR. Before machine-learning approaches were available, several computational methods were developed to search for other possible G4-motifs, such as G4Hunter ^15^, PQSfinder ^16^ This method algorithmically detects divergent G4 sequences that differ both in loop and tetrad conformation. Machine-learning models on gradient boosting and trained on G4-seq data, called Quadron, were next implemented ^17^. CNN models, such as PENGUINN ^18^ and G4detector ^19^ followed and were trained on G4-seq in vitro data.

Another CNN model - DeepG4 ^20^ was trained on combined G4 ChIP-seq and G4-seq data and the genome-wide maps of open chromatin regions(DNase-seq and ATAC-seq). With this approach the authors could detect cell-type specific active G4 regions, and identify key TFs predictive of G4 region activity. When combined, the methods predict nearly 1 million G-flipons in the human genome.

ENdb is an attempt to combine all different experimental G4 detection methods (G4-seq, G4 ChIP-seq and G4 CUT&Tag) methods. G4s in each dataset was called using pqsfinder with ≥1 bp overlap with G4 peaks required and the locus referred to as a eG4 (endogenus G4). A level of confidence to each eG4 from 1 to 6 based on how many tissue datasets the putative G4 fold was found. Of 391,503 eG4, 123,150 (31.5%) were found in only one tissue. The highest level was set at 6 and correspond to 27,376 (7%) eG4s detected in more than 10 samples regardless of the detection method ^21^.

### Large Language Model in Genomics

Large language models are the next generation predictive models that outperformed CNNs in many areas due to their ability to learn context. We previously applied DNABERT to reveal Z-DNA forming potential in human and mouse genomes ^22^, and here we apply DNABERT to predict G4s. As the training set, we use Kouzine et al data set generated with permanganate/S1 Nuclease Footprinting methods ^11^. This method has some advantages over ChIP-seq and CUT&Tag methods. It was applied to live cells with 70 sec exposure to potassium permanganate. This allowed it to catch non-B DNA structures that have unpaired thymines at junctions between B-DNA and an alternative conformation, such as in G4 loops and that are transiently exposed as the flip from one conformation to the other occurs. Since it takes energy to initiate the change in flipon conformation, these regions correspond to functionally important segments of the genome, as it will be seen from our analysis. The mapping approach has higher resolution than G4 ChIP-seq and G4 CUT&Tag methods, and provides predictions at base resolution that are not motif-based like those made by PQSfinder. The method minimizes problems associated with permeabilization of cells and long incubation periods in the presence of G4 stabilizing reagents. However, just as these other methods are limited by the preference of reagents such as BG4 for a subset of G4s, the training set is limited by the need for thymines that are reactive with permanganate in G-flipons.

## Results

### G4-DNABERT model

G4-DNABERT model was created by fine-tuning DNABERT ^23^ model trained on 6-mers representation of DNA sequence and used 512 bp context length (see details in the Methods). We used KEx ^11^ experimental data set for training (around 53 000 G4s) as it has higher than ChIP-seq resolution and was done on intact cells with 70 sec exposure to reagent. This method captures transient formation of G4 structures in ssDNA flipping from one conformation to another, or to thymines present in G4 loop sequences. With G4-DNABERT model we recalled 99% of KEx data set and predicted 3.4 times more G4s comprising a 180,000 G4s dataset (Supplementary Table 1). Distribution of G4-DNABERT G4s and KEx G4s over genomic regions (Supplementary Figure 1) revealed novel G4s in promoter regions (65.7% in KEx and 46.1% in G4-DNABERT) and a considerable proportion in non-coding regions - 25.6% over 15.2 % in intergenic and 25.6% over 14% in introns.

The main advantage of transformer-based model is that they can learn the context. The high attention scores reveal sequences that the model paid more attention to while learning genomic regions of interest. Attention score maps for G4 regions for MYC and KRAS promoters are presented in Figure 1C-E. The region around MYC promoter and G4s detected by different methods are presented in Figure 1C. Attention maps for the famous MYC G4 P1 promoter ^24, 25, 26^ is given in Figure 1D. The attention maps show that high attention corresponds not only to G4 regions but also to the immediate surrounding regions. The region around KRAS promoter ^27, 28^ and G4s detected by different methods are presented in Figure 1D. Attention maps around two G4s in KRAS promoter are given in Figure 1E. There are high attention scores at the boundaries of G4 region. This property of language models to learn context of functional regions allows for more refine selection of seemingly simple pattern. Analysis of 20 bp flanking regions around G4-DNABERT predicted regions revealed an extra G or C-repeat pattern at both sides from G4. This finding is line with observation that functional G4s such as in MYC or KRAS have extra GGG/CCC blocks, a so called “spare-tire” to ensure robust formation of G4 in the face of DNA damage ^29^. All 6-mers ranked by attention scores with corresponding frequencies in the genome are given in Supplementary Table 2.

**Figure 1.**
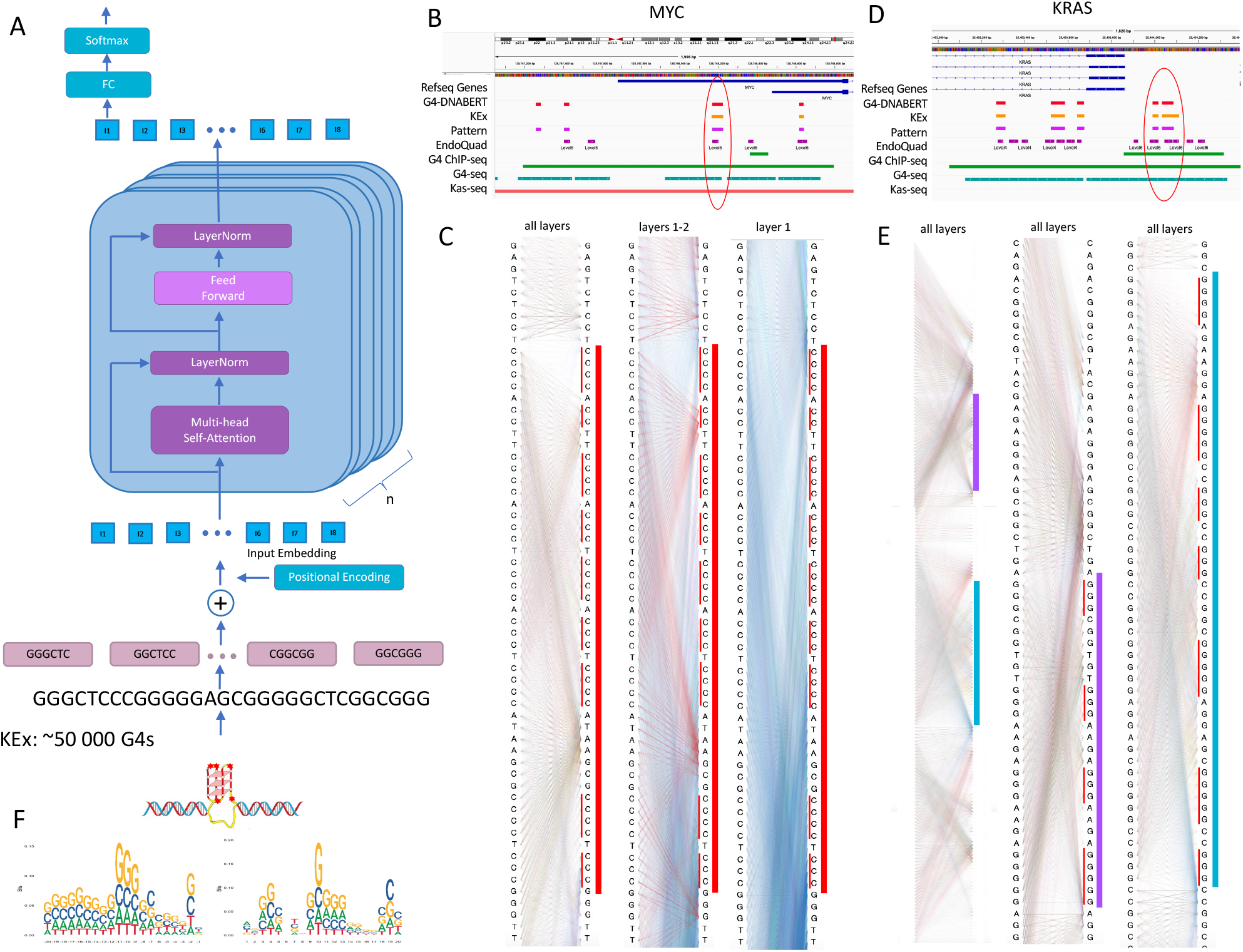
G4-DNABERT model. **A**. Architecture of G4-DNABERT model. It was made by fine-tuning DNABERT on KEx data. **B**. Quadruplexes at MYC promoters predicted by different methods. **C**. Attention maps of the region around G4 in MYC promoter encircled in red in Figure 1B. Left columns is the summary view from all 12 attention layers; middle column is the combination of layer 1 and 2; right column is an attention map from layer 1. Red bar indicates G4. **D**. Quadruplexes at KRAS promoters predicted by different methods. **E**. Attention maps of the region with two G4s at KRAS promoter (encircled in red in Figure 2D). Left column depicts the summary view of all 12 attention layers; middle column is the attention map from all layers around the first G4; right column is the attention map from all layers around the second G4. Violet and light blue bars indicate G4s. **F**. Logo of 20 bp flanks around G4s predicted by G4-DNABERT.

### G4-DNABERT model revealed novel G4s in non-coding regulatory regions

Comparison of G4-DNABERT with other G4 detection methods such as G4-seq, G4-ChIP-seq, KAS-seq, G4 Cut&Tag, EndoQuad database and computational pattern search is given in Figure 2A. The results from pairwise overlaps of eight different methods confirms the poor and varying overlap from 3% to 97%.

**Figure 2.**
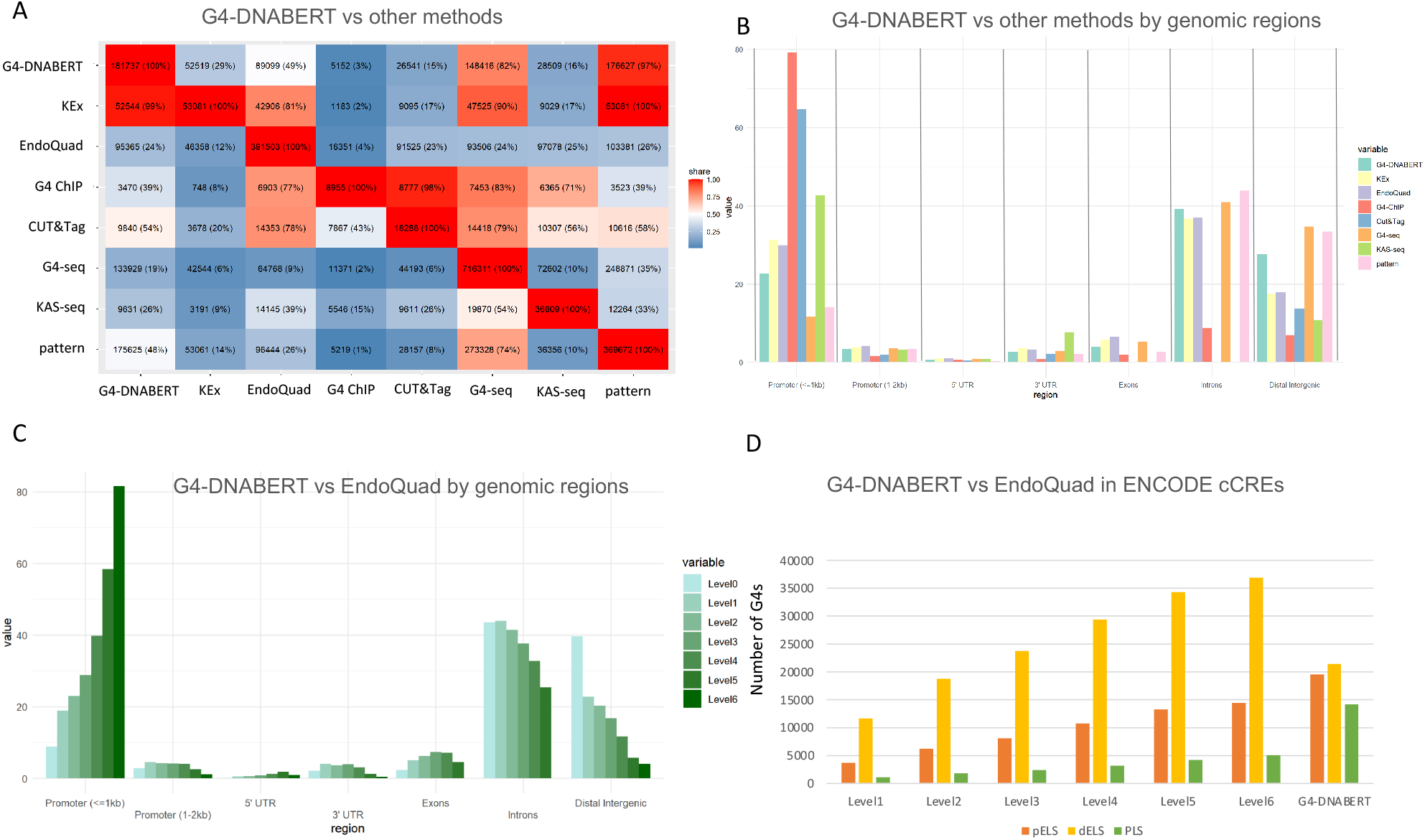
G4-DNABERT comparison with other methods. **A**. Genome-wide comparison of G4-DNABERT and other experimental G4-detection methods. **B**. Comparison of G4-DNABERT with other experimental G4-detection methods over genomic regions. **C**. Comparison of G4-DNABERT with EndoQuad database stratified by the confidence levels. **D**. Distribution of G4-DNABERT and EndoQuad over ENCODE cCREs.

EndoQuad contains today the most comprehensive set of G4s with corresponding levels of confidence ranging from 1 to 6 with level 1 being the lowest meaning that G4 was detected only in one sample and level 6 being the highest when G4 was detected in more than 10 samples. Only 49% of G4-DNABERT predictions overlap with the entire EndoQuad set (Figure 2B) confirming that there are still functional G4s to be experimentally mapped. G4-DNABERT and EndoQuad intersection stratified by levels are given in Supplementary Table 3.

Distribution of G4s predicted by different methods over genomic regions are given in Figure 2B. As expected G4 ChIP-seq and G4 CUT&Tag are enriched in promoters with almost 80% and 65% of predictions falling into these regions. G4-DNABERT, KEx, G4-seq and EndoQuad have almost equal share in introns – around 40%. G4-seq has the maximum share in intergenic regions (37%) followed by classical pattern and G4-DNABERT. G4 ChIP-seq and G4 CUT&Tag have small shares in non-coding regions.

Comparison of G4-DNABERT intersection with EndoQuad stratified by different levels of confidence over genomic regions is given in Figure 2C. Level 6 G4s common to G4-DNABERT and EndoQuad are mostly enriched in promoters. The inverse trend is observed for intronic and intergenic G4s when the common G4s appeared to have mostly level 1 support in EndoQuad. Many G4-DNABERT predictions that lack EndoQuad support are in non-coding regions and consequently will require further experimental validation.

### G4-DNABERT G4s in promoters-enhancer pairs

#### ENCODE cCRE

To further explore regulatory potential of G4s, we overlapped G4-DNABERT predictions with cCRE from the Encode project ^30^. 19,566 G4-DNABERT predictions intersect with proximal cCREs (8.6-fold enrichment, p<0.001, permutation test) and 21,429 (1.9-fold enrichment, p<0.001, permutation test) with distal enhancers. The overlap of G4-DNABERT and EndoQuad in promoters (PLS), proximal enhancers (pELS) and distal enhancers (dELS) is given in Figure 2D. For proximal enhancers G4-DNABERT predicted 40% more G4s than EndoQuad. The low overlap of G4-DNABERT predictions with those of EndoQuad in cCRE suggests that G4-DNABERT is revealing G4s in promoters and enhancers that are novel as they have not previously been annotated using older methods.

#### ENdb

We further studied in more details the distribution of G4s in promoter and enhancers. ENdb is manually-curated enhancer database ^31^ in which promoter enhancer interactions were experimentally verified. We took 271 experimentally validated enhancer-promoter pairs and found that 71 genes have G4 in enhancer (1 bp overlap), 107 have G4 in promoters and 35 enhancer-promoter pairs have G4s in both (Supplementary Table 4). The G4 forming sites are bound by many TFs such as NOTCH3, MYC, SOX2, TBX3 and others, that function in developmental processes and cell differentiation and bind to G-rich motifs (“System development” GO:0048731, FDR 1.23e-07; “Animal organ development” GO:0048513, FDR 3.24e-07; “Tissue “GO:0009888, FDR 1.09e06; “Regulation of cellular biosynthetic process” GO:0031326, FDR 1.35e06; “Regulation of cell differentiation” GO:0045595 FDR 2.30e-06). The full list of GO-enrichment categories are given in Supplementary Table 5. When we enlarge the region and allow G4s fall into +/-50-100 bp around enhancer we get increase the G4 count from 809 (340 in promoters and 469 in enhancers) to 828 G4s (357 in promoters and 471 in enhancers) but only generate two more G4s pairs, showing that the overlap of G4s at sites of cCRE are important for promoter-enhancer contacts.

#### Zoonomia conserved cCREs

One million cCREs conserved over 241 mammals were identified in the study of Zoonomia project ^32^. The total number of ELS regions was 809 429 pairs, from which we took the top 5% dELS and pELS (by phylop score, n = 40 477) and checked the overlap with G4-DNABERT predictions in a 2 kb region surrounding a promoter to identify G4 EP pairs (Supplementary Table 6-7). For 7 094 top 5% conserved pELS we identified 3 850 genes with G4pEP pairs (54%) and for 33 383 top 5% conserved dELS we identified 1 327 genes (1%) G4dEP pairs (Table 1).

**Table 1.** G4 enhancer-promoter (EP) pairs detected with different methods.

|  | **G4-DNABERT** | **sCell CUT&Tag** | **List of genes for G4 EP pairs / enrichment** |
| --- | --- | --- | --- |
| ENdb | 35 | 0 | STable 4 |
| Zoonomia conserved cCREs (top 5%) pELS | 3 850 genes | NA | STable 6 / STable 7 |
| Zoonomia conserved cCREs (top 5%) dELS | 1 327 genes | NA | STable 6 / STable 7 |
| osteosarcoma samples U2OS | 416 | 594 genes | STable 8 |
| breast cancer samples MCF7 | 4 | 79 genes | STable 8 |
| osteosarcoma U2OS + breast cancer MCF7 | 384 genes | 600 genes | STable 8 |

Enrichment analysis of these 3,850 G4 pEP pairs with G4s revealed the following GO-categories: “Protein binding” GO:0005515, adjusted p-value 5e-75; “transcription regulator activity” GO:0140110, adjusted p-value 9e-72, “sequence-specific DNA binding” GO:0043565, adjusted p-value 1e-67, “ulticellular organism development” GO:0007275, adjusted p-value 2e-123; GO:0007399 “Nervous system development”, adjusted p-value 2e-108. The full list of GO-enrichment categories is given in Supplementary Table 7. Of that list 500 of G4 pairs are found in EndoQuad DB with 3350 genes being unique for G4-DNABERT.

Intersection of G4-DNABERT G4s and the highest confident level of EndoQuad (Level 6) EP pairs includes many circadian rhythm genes (TEF -Thyrotroph embryonic factor, BHLHE40 - Class E basic helix-loop-helix protein 40, Transcriptional repressor, EGR1 - Early growth response protein 1; Transcriptional regulator. KLF10 - Krueppel-like factor 10; Transcriptional repressor, HNRNPU Heterogeneous nuclear ribonucleoprotein U; DNA- and RNA-binding protein; NCOA2 Nuclear receptor coactivator 2; Transcriptional coactivator for steroid receptors and nuclear receptors.)

#### sCell CUT&Tag G4s

Single cell G4 detection methods provide an opportunity to reveal functionally active G-flipons in one cell. We verified colocalization of G4s in EP pairs in single cell G4-CUT&Tag data for 5363 cells (4 samples with 2115, 2052, 352 and 844 cells) ^33^.

We tested G4pEP pairs (i.e. between pELS and PLS) using ENCODE cCREs tracks for proximal enhancers. For each promoter with G4 we searched for enhancers with G4 in a region of 2 kB. We totally detected 2384 transcripts of 600 genes with statistically significant enrichment in G4pEP pairs (FDR<0.05) (Supplementary Table 8). We performed the test on 4 different samples from 2 types of cells – osteosarcoma cells (U2OS) and breast cancer cells (MCF7). The important observation that was replicated in each sample is that the number of G4 enhancers were several times higher than the number of G4pEP pairs (Figure 3A). Out of 600 genes we identified 50 genes with more than 50% of G4pEP pairs relative to the number of G4 in enhancers. 42 G4pEP pair genes were unique for osteosarcoma cells and 8 were unique for breast cancer cells (the discrepancies reflect the different number of cells in different samples with osteosarcoma cells having around 2000 cells in each sample and breast cancer samples having 844 and 352 cells) (Supplementary Table 8).

**Figure 3.**
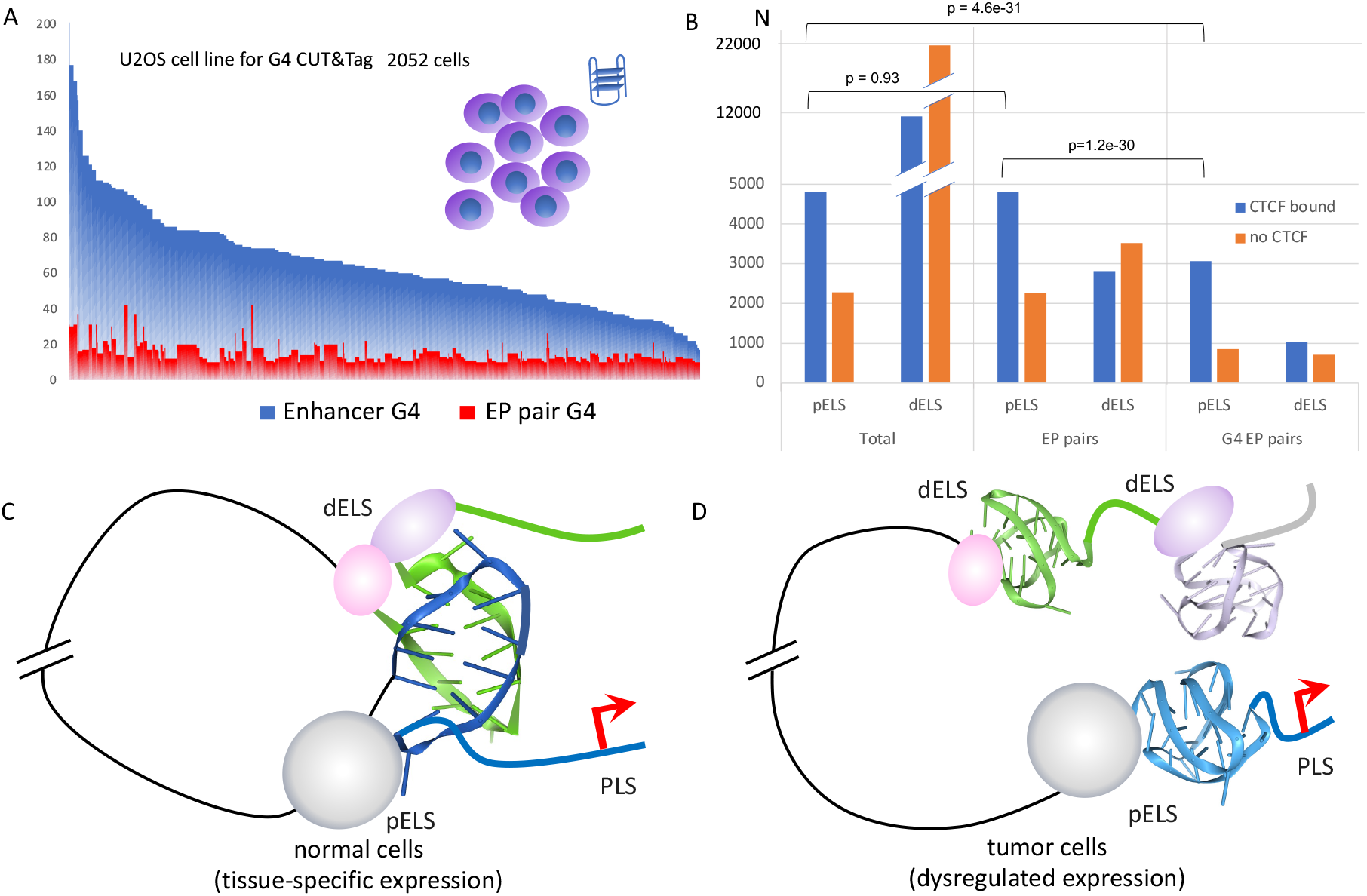
G4-DNABERT and EP interactions. **A**. Number of G4s in enhancers and in EP pairs for all genes that showed significant enrichment in G4 EP pairs in sCell G4 CUT&Tag experiment. Plot is given for one sample from U2OS cell line. Proximal enhancers are taken from ENCODE cCRE. **B**. CTCF colocalization with pELS, dELS and G4 EP pairs. **C**. Schematic representation of EP interactions by means of G4s. **D**. Poised enhancers with G4s waiting for the G4-binding proteins to initiate the transcription of the gene. Enhancer RNA (eRNA) can also form G4s and participate in interactions with promoters via R-loops.

We also tested how well G4pEP pairs predicted in one cell line can be replicated in another cell line. Thus, there are only 15 genes with G4-EP pairs that were replicated in all four samples (among them are transcription factors ZNF687, SP4, suppressor SUFU, transmembrane proteins SYNGR4 and TMEM143, and non-coding regulatory RNA genes.

There are 533 (out of 594, 90%) G4pEP pair genes that are unique for osteosarcoma cells and not replicated in breast lines. And there are 6 (out of 79, 8%) G4pEP pair genes in breast cancer cells that are not replicated in osteosarcoma cells (Supplementary Table 8). Analysis of molecular functions of genes with G4pEP pairs revealed enrichment in transcription factors including E2F-family, SP-family, zinc-finger family TFs and others (see full list in Supplementary Table 8.

We also identified genes with G4pEP pair genes that are overexpressed in osteosarcoma (IF1, CEP131, KCNC3, PRIM2, SAP130, ACTR6, REEP5, RHCE, FOXJ2, SLC17A5, CHRAC1, LRRC10B, VTI1B, USP42, UGP2, GSTZ1, ZC3H6, SGPL1) and breast cancer (PPM1D, BCAS3, USP32, PARD6B, TUBD1, RPS6KB1, BRIP1, TRIM37, ZNF217) (Supplementary Table 8).

The summary table with G4pEP pairs detected in different experimental data sets with corresponding number of genes and links to Supplementary tables is provided in Table 1. It is important to note that in estimating the number of G4pEP pairs in sCell CUT&Tag experiments, we underestimate the number of potential interactions due to the fact that CUT&Tag peaks are broad and may contain multiple G4s. Distribution of multiple peaks of G4-DNABERT and EndoQuad in CUT&Tag peaks for different levels of EndoQuad support for 4 cell lines are given in Supplementary Figure 1.

### CTCF and G4pEP pairs

We verified colocalization of G4s in proximal and distal enhancers and in G4pEP pairs and the results are presented in Figure 3B and Supplementary Table 9. We can see that G4pEP pairs are more often colocalized with CTCF, and the ratio for CTCF-bound pairs are higher for proximal enhancers. At the level of top 5% conserved enhancers from Zoonomia project, the proportion of G4pEP pairs reaches 80% for CTCF-bound enhancers (3068 out of 3919). The results point to the important role of G4s in regulation coupled with chromosome organization.

## Discussion

Based on the learned context G4-DNABERT replicate only half of all classical G4 sequence patterns creating a more refined set of regulatory G4s that is collated in EndoQuad. The collated collection in EndoQuad is based on experimental G4 detection whole-genome methods that often have poor overlap with each other where values vary from 1-2% as G4 ChIP-seq in classical pattern or in G4-seq datasets to 98% as CUT&Tag in G4-ChIP (Figure 2A). Each method has potential limitations. The G4-seq method is based on in vitro sequencing of DNA after extraction for cells and captures the difference in conformation when K^+^ or Li^+^ are used in sequencing buffers. The approach detected more than 700 000 G4s of which an unknown number may represent noise or G4 that are non-functional inside cells. ChIP-seq and CUT&Tag detect an order of magnitude less (around 10 000) but both depend on antibody that have preferences to G4s of particular topology (i.e. parallel orientation) ^2^. In our study with G4-DNABERT model we learned KEx data set (around 50 000) that mostly consists of classical G4s, but those that transiently formed in live cell, and extended this data set by two thirds at the genome-wide level. The advantage of G4-DNABERT is that it learns not only regular sequence pattern but implicit patterns in loops and implicit patterns of adjacent flanks as one can see in attention maps. A limitation of G4-DNABERT is the requirement for an unpaired thymines at the site of G4 formation. The analysis of G4-DNABERT 20 bp flanks revealed that often functional G4s have extra GGG/CCC-tracks as if to secure that G4 is formed when one block is broken. An excessive number of GGG or CCC repeats is observed in MYC (8 blocks) and KRAS (7 blocks) promoters. Taken in conjunction with those G4 listed in ENdb, the G4-DNABERT results help in the construction of a high confidence G-flipon set for further studies of the human genome.

G4-DNABERT revealed statistically significant enrichment of G4s in proximal (8.6-fold) and distal (1.9-fold) enhancers. We were curious whether G4s formed in these different regions could interact, so we searched for genes in which G4 elements were experimentally validated in both the enhancer and promoter regions. Analysis of experimentally confirmed EP pairs from ENdb or in conserved cCREs revealed almost 1500 EP pairs with G-flipons both in promoter and enhancer. For 7 094 top 5% conserved pELS, we identified 3 850 genes with G4pEP pairs (54%), consistent with about 50% EP pairs present on 25% of the same chromosomes assuming a random distribution. The result suggests that the formation of G4 in promoters and proximal enhancers is not linked, but likely reflects transcriptional processes that induce G4 formation at each site separately, as we will discuss.

We attempted to examine the issue further by using single cell sequencing datasets. However, the interpretation of that data is affected by the polyploidy of those cell lines. As documented by the American Type Collection, chromosome counts in the U2OS cells are in the hypertriploid range while those of MCF7 range from hypertriploidy to hypotetraploidy. By chance, 83% of cells would have generated experimental data for the joint presence of both enhancer and promoter G4 folds in each G4pEP pair (with only 67% present on the same chromosome, with the G4 formation by enhancers and promoters occurring on different chromosomes in other cases). We observed much less than this figure, suggesting that tumor cells have active mechanisms that diminish the frequency of G4pEP pair, likely through the unrestricted pairing of enhancers with alternative promotors, enabling the promiscuous transcription of many genes. The phenotypic diversity generated helps the tumor to evolve and survive. In this scenario, G4pEP pairs are important for the tissue-specific regulation of transcription and help enforce cell fate. The model is consistent with the recent report of frequent alternative promoter usage in tumors ^34^.

It is interesting to compare these findings with those reported for other data sets. Thus, the G4s revealed by CUT&Tag on mouse embryonic stem cells were localized at active promoters (characterized by the presence of eRNAs and both H3K4me1 and H3K27ac histone marks) and poised enhancers (H3K4me1 and of H3Kme3 marks), but not at primed enhancers (with only H3K4me1 marks) ^35^. Thus, the presence of G4 was proposed to be a functional structure that distinguishes active from primed enhancers in mouse embryonic stem cells .Importantly, the G4 structures existed independently of ongoing gene transcription. In the same study it was shown that G4s are lost at enhancers upon differentiation to neural progenitor cells, consistent with suppression of gene expression by heterochromatin formation.

In these studies, it was proposed that R-loops both initiate and maintain G4 formation. The model is supported by the genome-wide R-loop CUT&Tag experiments that revealed co-occurrence of single-stranded DNA, G4s and R loops at promoters and enhancers ^35^. In a separate study, the authors show that R-loops facilitate bunding of CTCF by G4 formation in many regions of mouse embryonic stem cells ^36^ and contribute to CCCTC-binding factor (CTCF) docking at cognate sequences. The CTCF mediated interaction contribute to long range interactions of genomic regions that cause looping to DNA to produce topologically associated domain (TAD) boundaries. Consistent with a role in chromatin organization due to effects on CCCTC binding, G4s are also enriched at TAD boundaries as demonstrated by the analysis of G4-seq and G4 ChIP-seq data sets ^37^. The authors further showed that adjacent boundaries containing G-quadruplexes frequently interact with each other contributed strongly to insulating the changes occurring within a particular TAD from those in adjacent regions.

These barriers then enable tissue-specific gene expression and prevent the spread of heterochromatin from adjacent regions. In another study ^37^ the authors demonstrated enrichment of G4s (based on G4-seq data) at chromatin domain boundaries and at CTCF binding sites. The CTCF anchors are asymmetrical placed at each end of the loop to confine cohesins within a TAD. Sliding of cohesin then promotes pairing within the TAD of distal enhancers with promoter regions (which by the author’s definition includes ENCODE proximal enhancers). In differentiated cells, the formation of loops promoted by G4 and CTCF-binding site underlies tissue-specific gene expression. In tumor cells, these stable loops are never formed, increasing the number of possible pairings between distal and proximal enhancers. The loss of regulation results in the use of alternative promoters rather than those that are tightly regulated in normal tissues and is consistent with the decreased formation of ENdb G4pEP pairs that we observe.

When taken together, the results from the different experiments support a role for G4 pairs in chromatin organization by bringing together distant elements. As our analysis reveals, the organization is less restrained in tumor cells. In contrast, differentiation of normal cells relies on the regulated selection of G4pEP pairs, as the mouse embryonic data suggests. The enrichment of G4 forming elements we observe near promoters suggests that these distant contacts help define and limit the promoters in use by a cell at a particular time during development.

RNA can also form G4s ^38^. The regulatory role of RNA G-flipons are still under investigation. It is possible that these long range G4 interactions are mediated by RNAs. A model, in which G4 in eRNA promote active transcription from enhancers has been proposed ^39^. In this scheme, G4-forming eRNA inhibits DNA methylation by sequestering PRC2, a repressive complex ^40^. The ibinding of G4 RNA with PRC2 causes dissociation of the enzyme from its nucleosome substrates, resulting in H3K27me3 depletion. The resulting change in chromatin structure enables gene activation. The model finds support from the study of G-flipons in a sheep model that revealed enrichment of G4s in regulatory regions similar to that found in other mammals ^41^. In sheep, 75% of the eRNAs have at least one G4. We cannot address such issues with G4-DNABERT, which is trained on KEx data and does not differentiate between DNA and RNA G4s.

## Conclusions

G4-DNABERT model, trained on KEx dataset, which was obtained from live cells and with technique that can obtain much narrow peaks compared to G4 ChIP-seq and G4 CUT&Tag methods helps further validate the formation of G4 structures in cells using an approach orthogonal to those documented in the ENdb database. A well curated G-flipon set now exists for further experimental exploration of roles for G4 in chromatin organization that incorporates our snapshot analysis of G4 formation in intact, living cells. G4 DNABERT also revealed many novel G4s in non-coding regions that are enriched in enhancers and promoters, with the more than 8-fold enrichment in ENCODE proximal enhancers. We verified G4 colocalization in enhancers and promoters in experimentally verified datasets such as ENdb, ENCODE cCREs, mammalian conserved cCREs from Zoonomia project, and sCell CUT&Tag 4 experiments. When considered with other findings, it is likely that G4 formation is involved in forming distant contacts with proximal enhancer regions that define and limit the set of genes expressed normally by a cell and the promoters used. Further experimental validation is required to delineate whether it is G4 formed by DNA or by RNA that program cellular differentiation by restricting TAD formation.

## Methods

### G4-DNABERT model

To create G4-DNABERT, we utilized a DNABERT model that had been pre-trained for 6-mer sequences (as detailed in ^23^). This model was specifically adapted, or “fine-tuned”, using KEx data ^11^. The fine-tuning process extended over 10 epochs, employing the ADAM optimizer. We set the maximum learning rate at 1 × 10−5 and chose a batch size of 24. During the initial 3 epochs, the learning rate incrementally increased from zero to its maximum value. For the subsequent 7 epochs, it gradually decreased back to zero.

Similar to our earlier model, Z-DNABERT ^22^, we developed five different models. Each was trained on 80% of the available positive class examples, along with a random selection of negative class examples. We made predictions for each 512 bp region of the entire genome by averaging the outputs of models that hadn’t encountered the data in their training phase. Thus, the final predictions represented an average from an ensemble of 5 models for new data and 4 models for previously used training data, including all experimentally identified positive regions.

Our experiments indicated that the accuracy of our predictions heavily relied on the context of the data. Consequently, for predictions encompassing the whole genome, we only used the central 128 base pairs from each 512 bp region. These pairs are surrounded by a minimum of 192 base pairs on either side, providing ample context. We achieved complete genome coverage by systematically shifting the starting point of each region by 192 base pairs.

### G4 experimental data

G4 experimental data sets used in this study were taken from the original publications: KEx ^11^, G4-seq ^9^, G4-ChIP ^42^, G4 CUT&Tag ^13^, KAS-seq ^43^, EndoQuad ^21^. Single cell G4-CUT&Tag data were taken from ^33^.

### Enhancer-promoter data

ENCOCE cCREs were donwloaded from UCSC genome browser. ENdb promoter-enhancer pairs were downloaded from the site www.licpathway.net/ENdb ^31^. Zoonomia conserved cCREs were taken from Zoonomia project ^32^.

### Data reprocessing and analysis

KEx raw data was downloaded from SRA (SRA072844). Quality control of KEx reads was performed using FastQC ^44^, reads were trimmed and aligned to GCRhg38. Deeptools computeMatrix and plotHeatmap were used to plot Figures. Random regions of the same size were used as control.

Cut&Tag raw data were downloaded from SRA (SRP331010) and processed using the count function of the 10X Genomics Cell Ranger ATAC software package. Unique fragments were used to plot TSS and UTR profiles. Deeptools computematrix and plotHeatmap were used to plot Figures. Conservative cCRE annotation was taken from paper ^32^.

Sequence LOGO’s were calculated and drawn using 20bp long flanking sequences of G4-DNABERT predictions via ggseqlogo R library.

Heatmap was build using published KEx, KAS-seq, G4-seq, G4-ChIP, Cut&Tag data. Pattern search was conducted using pattern (G{3,}[ATGC]{1,7}){3,}G{3} for each strand by a custom R script.

Scores for repeat elements were calculated using RepeatMasker annotation downloaded from UCSC.

Genomic distribution of tracks was calculated using ChIPseeker R library and UCSC ensGenes annotation.

## Data and Code Availability

Whole-genome annotation with G4-DNABERT is available as Supplemental Data 1. The code for G4-DNABERT model is freely available at https://github.com/mitiau/G-DNABERT.

## Contributions

DU developed G4-DNABERT, DK performed data analyses under the direction of AH and MP. AH and MP wrote the manuscript and prepared figures with assistance from DU and DK.

