## Supplemental Figure 1 for "Analysis of live cell data with G-DNABERT supports a role for G-quadruplexes in chromatin looping"

A

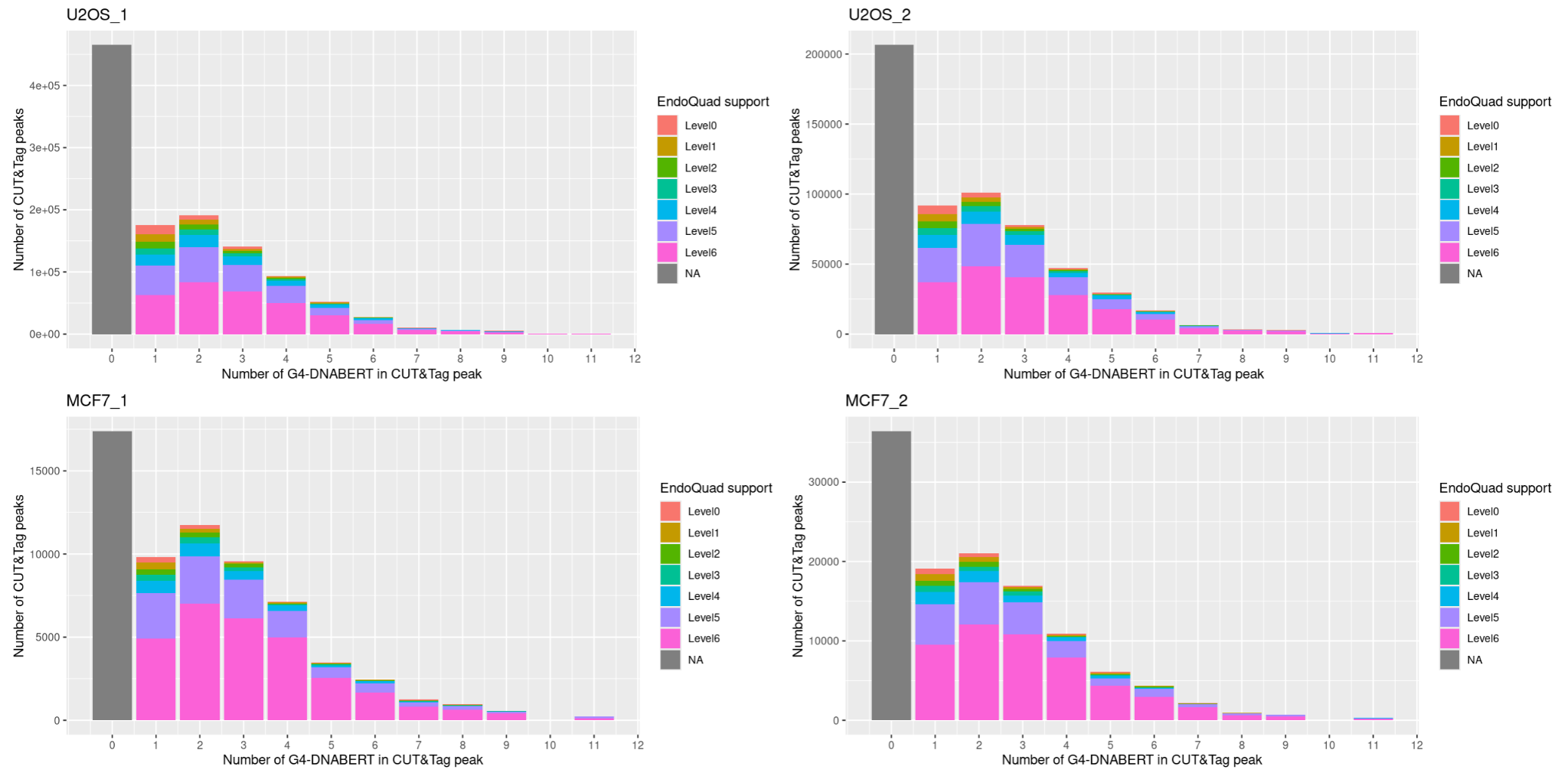

B

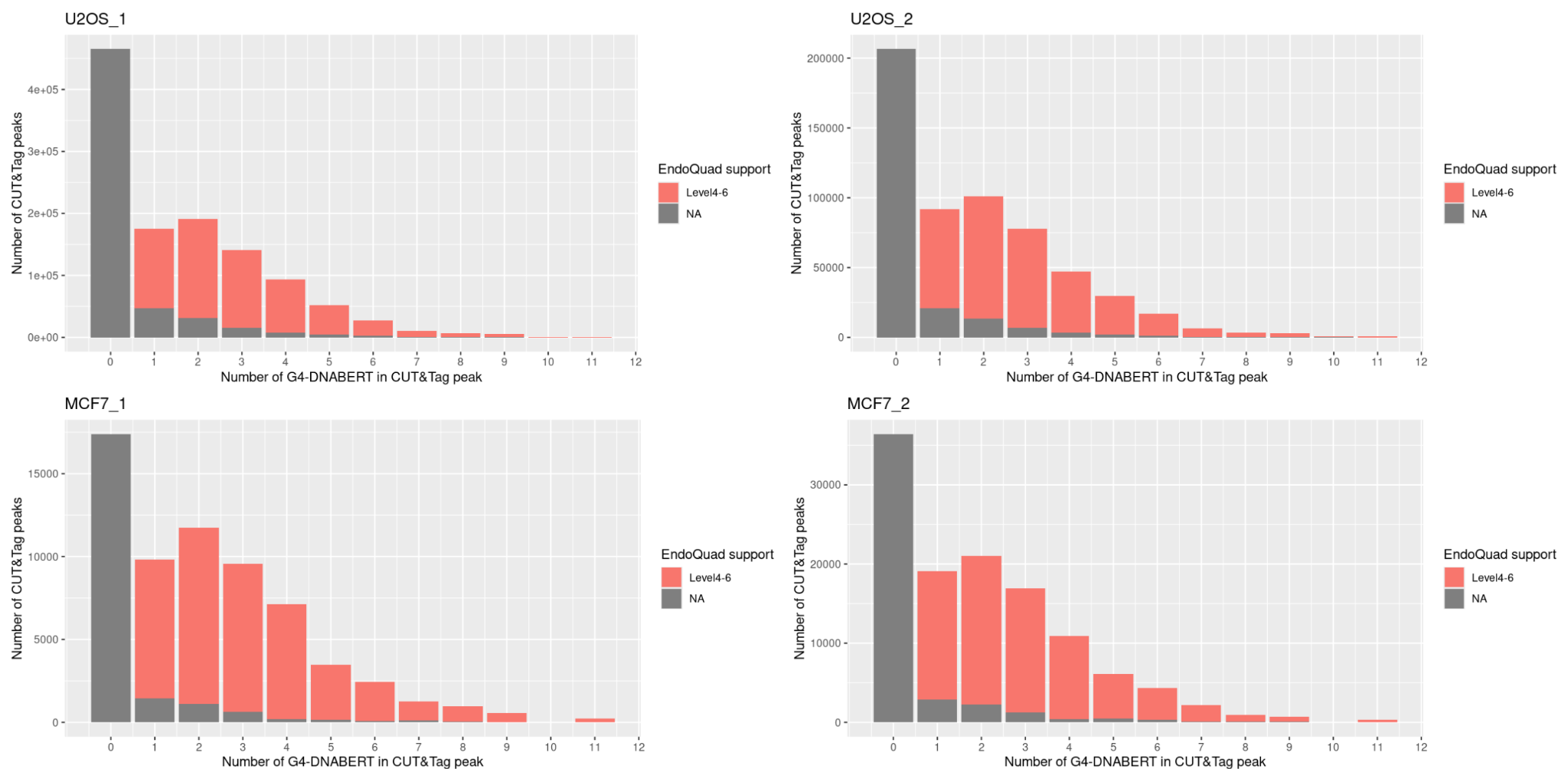

**Supplementary Figure 1.** Multiple peaks of G4-DNABERT and EndoQuad in CUT&Tag peaks. Different level of EndoQuad support is considered. **A.** All EndoQuad, levels 1-6. **B.** High support level of EndoQuad 4-6. NA means that the Endoquad falls below this threshold.
